# CAbiNet: Joint visualization of cells and genes based on a gene-cell graph

**DOI:** 10.1101/2022.12.20.521232

**Authors:** Yan Zhao, Clemens Kohl, Daniel Rosebrock, Qinan Hu, Yuhui Hu, Martin Vingron

## Abstract

In routine single-cell RNA-sequencing (scRNA-seq) analysis workflows, cells are commonly visualized in 2D to show the patterns in the data. However, these visualization approaches do not give any information about the genes that define the cell groups or clusters. It is therefore desirable to display cells and genes simultaneously such that by their relative position to each other information about the genes’ expression in a cluster can be obtained. Here we propose “Correspondence Analysis based Biclustering on Networks” (CAbiNet) as a novel approach to jointly visualize cells and genes by a non-linear embedding approach, called biMAP. The biMAP allows for easy and interactive exploration of cells and their corresponding marker genes in a single plot. CabiNet additionally offers an intuitive way to perform biclustering jointly on cells and genes, providing a simplified workflow to annotate cell types on the biMAP. CAbiNet is accessible through GitHub as an R package.

## 1 Introduction

Over the past decade, vast amounts of single-cell RNA sequencing (scRNA-seq) data has been generated, covering data from different species, tissues, developmental stages and sequencing protocols. This poses new challenges to data analysis as well as computational method development. Consequently, scRNA-seq analysis suites such as Seurat (Hao et al, 2021), Monocle (Cao et al, 2019a) or Scanpy (Wolf et al, 2018) have been developed to facilitate analysis. Some common workflows that are implemented in most toolkits include pre-processing, feature selection, dimensionality reduction, clustering, differential gene expression analysis and cell-type annotation by detected marker genes. While these methods already allow researchers to study the heterogeneity among cells, there is room for improvement with respect to intuitive and interactive exploration of the data.

The search for cell types and marker genes is usually proceeded by clustering the cells, visualizing cell clusters in a 2D embedding with t-SNE (van der Maaten and Hinton, 2008) or UMAP (McInnes et al, 2020), and, finally, determining differentially expressed genes by, e.g., DESeq2 (Love et al, 2014). Users generally cross-reference a marker gene list with the clustering results to adjust parameters and annotate the cell types. However, this process is laborious, which has motivated us to simplify it by simultaneously visualizing cells and their marker genes in a single planar embedding.

A joint visualization of cells and genes is beneficial in several ways: firstly, it immediately shows how specific a gene is to a cell cluster and provides information on the cells and corresponding marker genes, making cell type annotation a more intuitive process. Secondly, the relative position of genes to cell clusters also shows how much of the overall expression of a gene is shared between clusters.

A principal component analysis (PCA) biplot can display cells and genes in the form of their principal component values and loadings, the latter being the weight of a gene as part of a principal component (PC). However, the cells’ PC scores and the genes’ loadings get displayed on different scales, making interpretation difficult. The correspondence analysis (CA) biplot (Greenacre, 2017) improves on this by re-scaling cell and gene coordinates and showing them in the same space, thereby making the cell-gene relationships more interpretable than in a PCA biplot. However, it is common to most linear methods that for large and complex data sets, a low-dimensional projection discards a lot of information. For example, in a single-cell transcriptomic data set the first two PCs often only explain 1-2 % of the variance/inertia in the data. Therefore, a large number of dimensions have to be kept to adequately represent the data, obviously at the expense of visual interpretability.

Nonlinear embedding approaches like t-SNE and UMAP visualize high dimensional cell coordinates in a 2D map. However, only few algorithms have been developed that allow a joint visualization of cells and genes in 2D space. Recently, Chen *et. al* put forward SIMBA (Chen et al, 2022b), a graph embedding tool that simultaneously embeds cells and features based on a multi-relational graph. This however requires discretization of the data and the distance between two entities in the embedded space can be difficult to interpret. Also, SIMBA is not readily integratable into other established analysis workflows.

Here we present “Correspondence Analysis based Biclustering on Networks” (CAbiNet) to produce a joint visualization and co-clustering of cells and genes in a planar embedding. CAbiNet employs CA to build a graph in which the nodes are comprised of both cells and genes. Then a clustering algorithm determines the cell-gene clusters from the graph. Finally, the cells, genes and the clustering results are visualized in a 2D-embedding. We call such an embedding a biMAP. Due to the geometry of correspondence analysis and cell-gene graph construction, the biMAP displays a cell cluster’s marker genes within or near the cell cluster, allowing for easy interpretation. Cells and genes from the same cluster are colored identically in the biMAP. CAbiNet is implemented as an R package and freely available to download from GitHub (https://github.com/VingronLab/CAbiNet). It is fully compatible with popular scRNA-seq analysis pipelines such as those from Bioconductor. We will give an outline of the CAbiNet algorithm with the full details presented in the Methods. Its utility for faithfully embedding cells and genes into 2D will be demonstrated on four different scRNA-seq and spatial transcriptomic data sets. We will show how the CAbiNet-generated biMAP can be used to accelerate cell type annotation and discover cell-types. The examples cover both small data sets with clearly delineated cell types and highly complex developmental data sets that demonstrate the ability of biMAP to produce effective visualizations even for intricate biological experiments. In order to assess the quality of the biclustering, we comprehensively benchmarked CAbiNet in comparison to other biclustering algorithms on simulated as well as on expert annotated experimental data. Additionally, we compared it to common cell clustering analysis workflows.

## 2 Results

### 2.1 Visualization and Biclustering of scRNA-seq data

CAbiNet takes a gene expression matrix as input, in which genes are presented in rows and cells in columns. The algorithm builds on the transformation of the data into a matrix of Pearson residuals as done in correspondence analysis (CA). CA decomposes this matrix into a singular vector matrix of cells (matrix **U** in Fig. 1, see also in Methods 4.2), a diagonal matrix with singular values and a singular vector matrix of genes (matrix **V** in Fig. 1). Singular vectors are sorted by importance - “inertia” in CA - which allows for projecting the data into a lower dimensional space with the highest inertias. Notably, CA provides a scaling of **U** and **V** so as to overlay them in the same space. This gives rise to interpretable cell-cell and gene-gene distances and makes cell-gene associations possible (Methods 4.2). Thus, we can create a k-nearest-neighbor graph connecting both genes and cells (see Fig. 1). This graph, in turn, is used to calculate the overlap among neighborhoods of nodes to generate a shared-nearest-neighbor (SNN) graph.

**Fig. 1.**
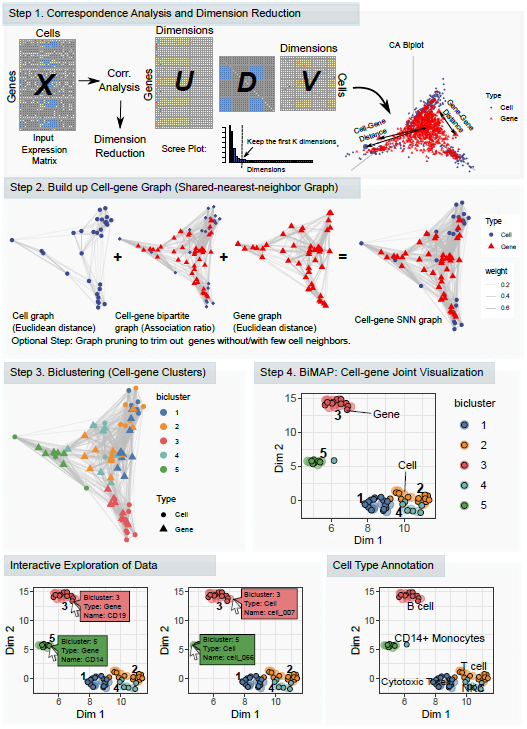
Overview of the CAbiNet algorithm. Step 1: Following correspondence analysis practice, the gene expression matrix **X** gets normalized and converted into a matrix of Pearson residuals, which is then decomposed by singular value decomposition into the left (**U**) and right (**V**) singular vector matrices. Step 2: kNN-graphs are built from a dimension reduced space based on either the euclidean distance, or, for the cell-gene/gene-cell graph, based on the association ratio (see Methods 4.3). The subgraphs are subsequently merged to form a single graph containing both cells and genes. If necessary, this graph is then pruned in order to remove spurious edges and converted to a shared-nearest-neighbor-graph (SNN-graph). The SNN-graph is the basis of both the biclustering and the biMAP. Step 3: Detect cell-gene biclusters from the graph. Step 4: biMAP visualization which displays the biclustering results with both cells and genes. Note that the biMAP can be plotted before biclustering to give users an intuition of how many biclusters are in the data. CAbiNet allows for an interactive exploration of the data. Hovering the mouse cursor over a point displays relevant information such as the cell/gene name and the bicluster. The biMAP intuitively shows the detected marker genes of each bicluster and enables a quick annotation of biclusters. For example *CD19* in bicluster 3 indicates that this bicluster consists of B cells and their marker genes.

The SNN graph serves as input to an embedding algorithm like UMAP or t-SNE to produce the biMAP. Since the SNN graph comprises both cells and genes, the embedding of the graph yields the joint visualization of cells and genes. The shared nearest neighborhood has been shown to be a good measure of similarity in high dimensional space such as gene expression data (Houle et al, 2010). Genes that are specifically highly expressed in a group of cells will therefore gravitate towards this cluster, while genes with constant expression profiles will be located close to the center of the embedding. By highlighting cell type marker genes, the biMAP can be used to quickly and easily annotate cell clusters or to identify marker genes through interactive exploration of the data.

The cell-gene graph also immediately suggests an intuitive strategy for biclustering: established clustering algorithms for large networks such as Leiden (Traag et al, 2019) or Spectral clustering (Donath and Hoffman, 1973; Fiedler, 1973; Von Luxburg, 2007) can co-cluster cells and genes in the SNN graph. A cluster can then contain both cells and genes, where, due to the design of the graph, genes that co-cluster with a set of cells tend to be more highly expressed in the cells they cluster with compared to other cells. We call those genes “associated to” or “specific for” the cells in the same cluster. The co-clusters can easily be understood as cell clusters with their corresponding marker genes. Conveniently, we can further adopt the concept of *S*_*α*_-scores from Association Plots (Gralinska et al, 2022) to rank genes by how specific they are for a cluster.

CAbiNet is implemented as an R package and can be installed from GitHub (https://github.com/VingronLab/CAbiNet). CAbiNet functions can replace corresponding procedures in routine scRNA-seq analysis piplines and are compatible with Bioconductor’s SingleCellExperiment object. The package will produce a biMAP with cells and genes colored according to the biclustering result. CAbiNet also allows for highlighting genes of interest in either a static or interactive biMAP. This promotes an intuitive exploration of genes and cells and facilitates cell type annotation. The Online Supplementary Material of this paper contains interactive html-files of biMAPs for the data sets discussed below. Users can mouse over the points in the biMAP to see the information of items including types (cell/gene), names and assigned biclusters.

### 2.2 Analysing scRNA-seq data with CAbiNet: PBMC10x data

We demonstrate the basic functionality of CAbiNet with a single-cell PBMC10x RNA-seq data set (Ding et al, 2020a). This data set comprises 3,176 cells annotated into nine cell types with 11,881 expressed genes detected. For our purpose, the advantage of this data set is that the authors have provided expert annotation of cells based on FACS. Among others, the nine annotated cell types include B cells, CD14^+^ monocytes, and natural killer cells.

As described above, CAbiNet performs dimensionality reduction, clustering and visualization. After standard pre-processing (Methods 4.1), CAbiNet computes CA, projects the PBMC10x data into a lower dimensional space and builds the SNN graph. CAbiNet then detects the biclusters and visualizes the results in a biMAP (Fig. 2a) by applying UMAP on the SNN graph.

**Fig. 2.**
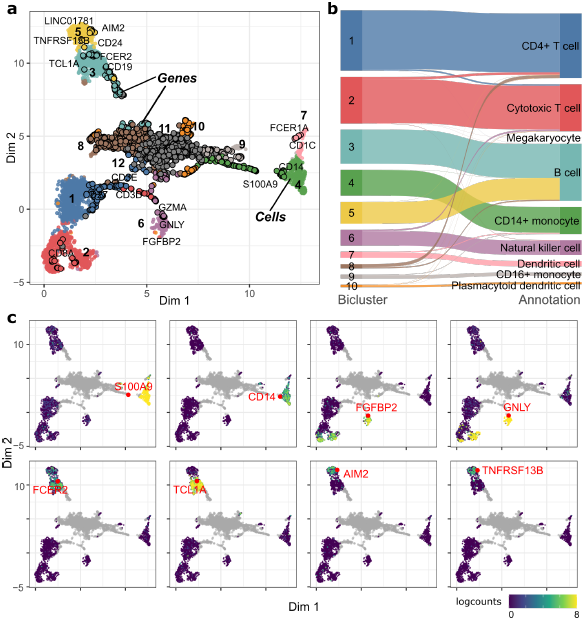
Application of CAbiNet on PBMC10x data. **a**, Joint biMAP visualization of the cell-gene biclustering results by CAbiNet, with genes and cells from the same bicluster colored identically. Genes are black circles filled in with the color of the associated cell cluster and cells are smaller dots. Some known marker genes have been labeled manually. An interactive version of this figure can be found in the Supplementary Data. **b**, The agreement between the expert annotation and CAbiNet biclustering results is shown in the Sankey plot. **c**, The expression levels and position of selected marker genes are shown on the biMAP. The grey points are genes and cells are colored by the log_2_-expression levels of genes highlighted in red. CD14^+^ monocytes marker genes *S100A9* and *CD14* in bicluster 4 are highly expressed in cells that co-clustered with them. The natural killer cells marker genes *FGFBP2* and *GNLY* are highly expressed in the co-clustered cells in bicluster 6. *FCER2* and *TCL1A* are highly expressed in bicluster 3, while *AIM2* and *TNFRSF13B* are highly expressed in bicluster 5, indicating that cells in these two clusters are different B cell subtypes.

The clustering quality is commonly measured by the adjusted rand index (ARI, see Methods 4.14). CAbiNet achieves an ARI of 0.79 on this data set, indicating a good agreement between the CAbiNet clustering and the expert annotation. Fig. 2b shows a Sankey plot illustrating the correspondence between annotation and computed clusters. The large agreement allows us to compare our results with the expert annotations.

The biMAP in Fig. 2a shows clusters of cells and genes, the latter represented by black circles filled in with the color of the associated cell cluster. The clusters located in the center of the biMAP (clusters 11 and 12) are composed exclusively of genes, which are not specific to any cell cluster SFig. 1 shows that these genes are ubiquitously expressed among cell types. As such they do not contribute information towards cluster annotation.

The biMAP places cell-type specific genes close to the corresponding cell groups. We manually labeled known marker genes for cell types to allow for easier interpretation and validation. For example, *S100A9* and *CD14* are located towards cluster 4 (Fig. 2a) and are known marker genes for CD14^+^ Monocytes, immediately suggesting that cluster 4 corresponds to this cell type. The feature plots in Fig. 2c confirm that indeed these two marker genes are highly expressed in this cluster. The Sankey plot in Fig. 2b also confirms the identity of cluster 4 as CD14^+^ monocytes. Likewise, the natural killer cell marker genes *FGFBP2* and *GNLY* are close to cluster 6, suggesting the identity for this cell cluster, which is also supported by their expression pattern in the feature plots (Fig. 2c) and by the mapping to the expert annotation in the Sankey plot (Fig. 2b).

Interestingly, the biMAP suggests a separation among the expert annotated B cells into two groups represented by the two clusters numbered 3 and 5 in Fig. 2a-b. Each of the two subgroups has its own set of marker genes. Note that the color code for the genes coincides with that of the cells from the same cluster. Cluster 3 in cyan-blue appears to be naive B cells based on the proximity to naive B cell marker genes *FCER2* and *TCL1A* (Fig. 2a) (Ramesh et al, 2020). Likewise, genes *AIM2* and *TNFRSF13B* belong to cluster 5 (light-yellow color) and their association with memory B cells (Ramesh et al, 2020; Franzén et al, 2019) suggests this identity for cluster 5 (Fig. 2a). The gene expression levels shown in the feature plot in Fig. 2c support this interpretation.

### 2.3 Tabula Muris Limb Muscle data set

We further apply CAbiNet to the Tabula Muris Limb Muscle (abbrv.: tabula muris) data set (Schaum et al, 2018). The data set consists of 1,960 cells and 20,214 genes and was annotated by experts using cell-type specific marker genes to label the clusters. After running CAbiNet, the data is clustered into eight clusters of both cells and genes (Fig. 3a). Each of the biclusters consists of several hundred cells and a similar number of genes. Cluster 8 in the center of the embedding consists of genes that are non-specifically expressed among clusters. The two cells co-clustered with these genes coincidentally express a large number of these non-specific genes. The Sankey plot in Fig. 3b shows that the cell clusters show good agreement with the expert annotation (ARI of 0.75).

**Fig. 3.**
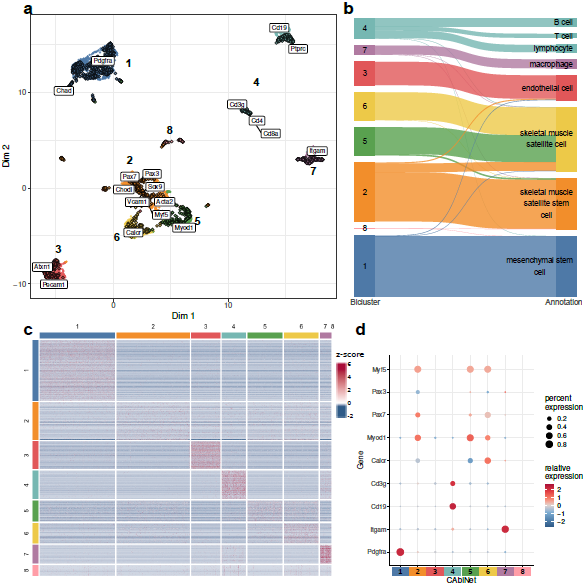
Analysis of Tabula Muris Limb Muscle data with CAbiNet. **a**, biMAP of the Tabula Muris Limb Muscle data with marker genes identified in the original publication and those discussed in the main text labelled. An interactive version of the biMAP is available in the Supplementary Data. **b**, Sankey plot showing how the expert annotation corresponds to the CAbiNet cell clusters. Cell types are colored by the cluster with the greatest contribution to it. **c**, The heatmap shows the z-scores of the log-normalized expression of CAbiNet biclusters with cells in the columns and genes in the rows. **d** The expression of the genes discussed in Results 2.3 is shown in the dot plot.

Figure 3a shows how cell type specific marker genes (as defined in the original publication) are located at the relevant (sub-)clusters. Platelet-Derived Growth Factor Receptor Alpha (*Pdgfra*), a mesenchymal stem cell marker gene, is, as expected, embedded adjacent to the *Pdgfra* expressing cells in cluster 1 (SFig. 2a). Similarly, cells in cluster 7 consist of macrophages (Fig. 3b), and indeed the majority express the macrophage marker gene *Itgam*, which is located next to cluster 7 (SFig. 2b). T cells and B cells are clustered together (Fig. 3b) but can be immediately differentiated by *Cd19* (B cells) and *Cd3g* (T cells) expression (SFig. 2c-d). In this way, the biMAP offers an easy way to annotate and validate the clustering and can help to determine when a cluster should be divided.

Skeletal muscle satellite stem cells (cluster 2) and skeletal muscle satellite cells (clusters 5 and 6, Fig. 3b) have overall similar expression profiles, which can also be confirmed in the heatmap in Fig. 3c. This makes the assignment of a gene to one or the other cluster difficult. The original publication carefully analyzed these genes and observed that the satellite cells can be subdivided into two clusters based on the expression of *Myod1* and *Calcr* (clusters 5 and 6, SFig. 2e-f) depending on their activation status. In the biMAP, *Myod1* is located near cluster 5 while *Calcr* lies near cluster 6, in good agreement with the respective expression patterns and the observations from the original publication. Additionally, it can be observed from the biMAP that satellite stem cells (cluster 2) simultaneously express *Pax7* and *Pax3*, which can be confirmed in the respective feature plots (SFig. 2g-h) (Relaix et al, 2005). Genes that are equally shared between the three clusters (2, 3 and 5), such as *Myf5*, are located between them (SFig. 2i).

### 2.4 Cerebral Organoids data set

In order to test the performance of CAbiNet on a highly complex data set, we analyzed a scRNA-seq data set comprising 35,291 cells derived from human cerebral organoids (abbrv.: brain organoids) (Rosebrock et al, 2022). The data is of particular interest as it consists of fully differentiated neurons, undifferentiated neural stem cells, as well as cells in transitory states which can have a considerable overlap in their expression profiles. This data set thereby helps to highlight the capabilities and limitations of CAbiNet. For a detailed description of the data and the processing see Methods 4.9.

CAbiNet groups the organoids into 13 clusters (Fig. 4a). The Sankey plot in Fig. 4d shows that the biclusters overlap well with the 17 annotated cell types from the original publication (Rosebrock et al, 2022) (ARI: 0.59). As can be observed in Fig. 4a, the biMAP embedding divides the data into two major parts: The stem cells in the upper half, and more specialized cell types in the lower part. Additionally, cortical cell types are located on the right side while cell types towards the lower left belong to non-neural lineages. Additionally, a clear developmental trajectory can be observed from the cortical neural stem cells (NSCs) expressing *SOX9*, towards cortical neurons via intermediate progenitors (IPs) (Fig. 4a-b). Cluster 1, 6 and 7 consist of NSCs in different stages of the cell cycle (SFig. 4), with cells from cluster 7 comprised of dividing cells. Marker genes associated with differentiating neurons (*NEUROG2*) are situated in close proximity to cluster 4 (IPs), while markers for postmitotic cortical neurons (*NEUROD2, NEUROD6*) are embedded adjacent to the terminally differentiated cells (cluster 2) (Fig. 4a-b, SFig. 3a). Clusters 2, 3 and 4 have a large overlap in their expression profiles, but cells from cluster 3 co-cluster with genes more associated with regions of the midbrain/thalamus such as *FOXP2* and *TCF7L2* (SFig. 3b-c). Cluster 3 can be further subdivided by their co-clustered genes into two subpopulations: GABAergic inhibitory neurons (*LHX1, GAD1*) and excitatory glutamatergic neurons (*ADCYAP1, SLC17A6*, see SFig. 3d-g). The myelination factors *MPZ* and *EGR2* are embedded together with cluster 11, indicating that the cluster consists of Schwann cells (SFig. 3h-i). This is further confirmed by the co-embedding of *SOX10*, which has been shown to be required to maintain the Schwann cell identity (Finzsch et al, 2010)(SFig. 3j). Cells from cluster 13 are characterized by strong expression of *NEUROD1, PPP1R17, ISL1, POU4F1* and *SIX1*, all of which are embedded within the cluster (SFig. 3k-o). *NEUROD1, ISL1, POU4F1* (the gene producing BRN3A) and *SIX1* have been shown to play vital roles in sensory neuron development (Dykes et al, 2011; Deng et al, 2014; Sun et al, 2008; Sato et al, 2015) whereas *PPP1R17* is a typical Purkinje cell marker (Kozareva et al, 2021). This makes it difficult to assign a specific cell type to the cluster. Accordingly, the cells were labelled as “posterior CNS/PNS” cells in the original publication.

**Fig. 4.**
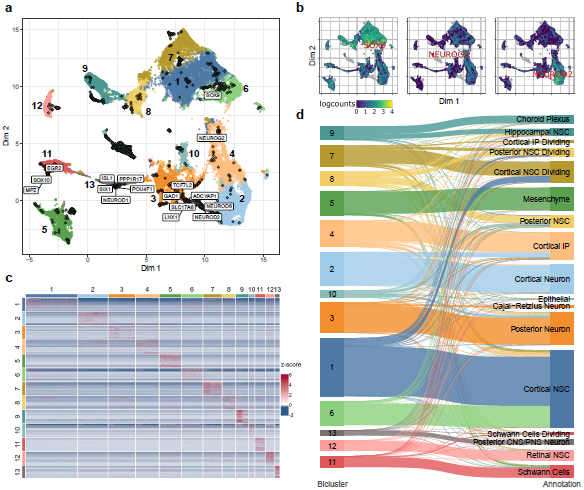
Analysis of brain organoid data with CAbiNet. **a**, biMAP of the brain organoid data set colored by CAbiNet biclusters. Marker genes discussed in the main text for specific brain regions and neuronal cell types are marked. An interactive version of the biMAP is available for download in the Supplementary Data. **b**, The feature plots highlight the placement of marker genes along the developmental trajectory from neural stem cell (*SOX9*) to IPs (*NEUROG2*) to fully differentiated neurons (*NEUROD2*). **c**, Heatmap showing the top 10 genes with the highest *Sα*-score per cluster. **d**, The Sankey plot illustrates how the cell clusters relate to the expert annotations. The cell types are colored by the cluster that contains the largest number of cells from the cell type.

As shown in the heatmap in Figure 4c, the top 10 co-clustered genes show distinct expression patterns and are consistently more highly expressed in their respective cluster. This shows that the biclustering indeed recovers relevant marker genes.

### 2.5 Spatial *Drosophila melanogaster* embryo data

Spatial transcriptomic data constitutes a particular challenge to the data analysis due to large number of drop-outs (Wang et al, 2022; Chen et al, 2022a). To further study the performance of CAbiNet on such sparse data, we applied it to spatial transcriptomic data of *Drosophila melanogaster* late-stage embryos (14-16h after egg laying (E14-16h)) (Wang et al, 2022). The gene expression profile was resolved by Stereo-seq (Chen et al, 2022a) yielding 14,808 pseudocells (bins of pixels on chip which are recognized as equivalent of cells by the original publication) with 7,178 genes. In the original publication 10 cell types were annotated based on unsupervised clustering. The standard UMAP projection shown in Fig. 5a illustrates that the boundaries between cell types are ill defined (e.g. epidermis vs. foregut and epidermis vs. trachea). This highlights the difficulty of distinguishing cell types and identifying corresponding marker genes in spatial scRNA-seq data.

**Fig. 5.**
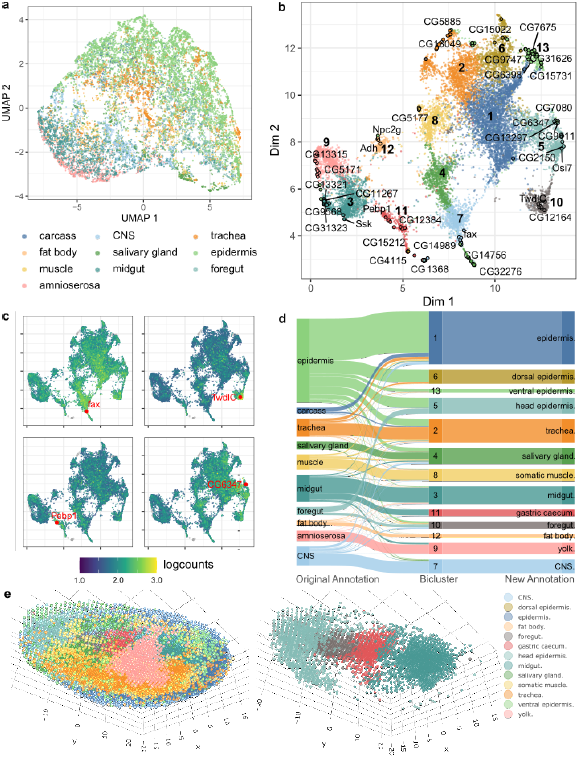
Spatial *Drosophila melanogaster* Stereo-seq data. **a**, UMAP embedding of expert annotated cell types. **b**, biMAP embedding of cell-gene biclusters. The genes and cells from the same biclusters are colored identically. Genes are filled circles with a black outline and the cells are the smaller dots. Selected marker genes are labeled in the biMAP. **c**, The feature-biMAPs show the expression levels of known marker genes (*fax* (CNS), *TwldC* (foregut), *CG6347* (head epidermis) and *Pebp1* (gastric caecum)) in the cells. The cells are colored by the log_2_-expression levels of the highlighted genes. **d**, The Sankey plot shows the consistency among the expert annotation, the biclustering results from CAbiNet and the revised CAbiNet-based annotations. **e**, Spatial distribution of the cells. The left panel is the 3D visualization of the embryo with cells colored by the biclustering. The right panel shows four cell types out of the left panel. From head to tail they are head epidermis, foregut, gastric caecum and midgut. Interactive versions of panel b and e can be found in the supplementary materials.

CAbiNet recognizes 13 biclusters from the data with biologically meaningful co-clustered marker genes. For example, 8 out of 14 genes (*fax, CG14989, Cam, Gbeta13F, Obp44a, ctp, fabp*) in cluster 7 are known marker genes of the central nervous system (CNS), and 6 out of 11 genes (*TwdlC, CG12164, Cpr50Cb, Cpr56F, Cpr65Av, Cpr66D*) in cluster 10 are known foregut marker genes. The expression levels of *fax* and *TwdlC* shown in Fig. 5c suggest that they are specifically highly expressed marker genes for the co-clustered cells.

CAbiNet also captures fine-grained cluster structure and offers an intuitive embedding of biclusters which can be used for cell type annotation. For example, we found that most cells annotated as midgut in the original publication using Scanpy correlate to two clusters by CAbiNet, cluster 3 and 11, which are recognizable as distinct groups in the biMAP (Fig. 5b,d). This indicates that cells originally annotated as midgut could be further divided into two cell types. Checking the detected marker genes in cluster 11, we found some of them, e.g. *Pebp1* and *Acbp4*, are known marker genes of gastric caecum. These genes have overall higher expression levels in cluster 11 compared to other clusters (Fig. 5c), indicating that cells in cluster 11 represent gastric caecum, a sub-structure of the midgut that was not previously identified in the original analysis. Similarly, we also found that cluster 5 represents head epidermis which is a subtype of epidermis. The expression level of head epidermis marker gene *CG6347* is shown in Fig. 5c.

Based on the biclustering results from CAbiNet, we generated new annotations of the cell clusters. The annotated cell types are shown in Fig. 5d. Figure 5e shows the cells in the embryo, color-coded by these enhanced annotations. Reminiscent of the actual embryonic anatomy, the spatial positions of annotated head epidermis, foregut, gastric caecum and midgut cells (right panel of Fig. 5e) are ordered from head to tail. Thus, CAbiNet provides a more informative joint embedding of genes and cells compared to Fig. 5a. CAbiNet also generates fine-grained biclustering results and supports cell type annotation for spatial transcriptomic data.

### 2.6 Evaluation on simulated data

In order to determine how well the biclustering performs on scRNA-seq data, we ran CAbiNet as well as eight other biclustering algorithms on simulated data generated using Splatter (Zappia et al, 2017) (see Methods 4.11). Splatter can learn and preserve the distribution patterns from real data and generate simulated scRNA-seq data which has distinguished cell clusters and corresponding differentially expressed genes. Since the clusters and their differentially expressed genes are known, one has a gold-standard biclustering to compare computational results to.

We evaluated the performance of the algorithms based on the Adjusted Rand Index (ARI) and the clustering error (CE) (Patrikainen and Meila, 2006; Padilha and Campello, 2017) (see Methods 4.14). The ARI provides a quantitative comparison between a computed cell clustering and a goldstandard cell clustering. The CE measures the quality of a biclustering. For both measurements a higher score indicates a better (bi-)clustering.

Simulated data are generated by Splatter in accordance with parameters derived from real data sets, which in our case are the Zeisel Brain Data (abbrv.: zeisel) (Zeisel et al, 2015) and the PBMC3k data from 10x Genomics (abbrv.: pbmc3k) (10x Genomics, 2016). Splatter also allows to determine the difficulty of clustering a data set via a factor on the mean expression strength on differentially expressed genes and by setting the frequency of differentially expressed genes (see Methods 4.11). We tested the CAbiNet implementation with both Leiden and Spectral clustering and compared to QUBIC (Li et al, 2009), s4vd (Sill et al, 2011), Plaid (Lazzeroni and Owen, 2002), Unibic (Wang et al, 2016), BiMax (Prelić et al, 2006), CCA (Cheng and Church, 2000), Xmotifs (Murali and Kasif, 2003) and IRIS-FGM (QUBIC2) (Xie et al, 2019; Chang et al, 2021). In order to better compare CAbiNet to popular scRNA-seq analysis workflows, we clustered cells and identified differentially expressed genes with two R packages, Monocle3 (Cao et al, 2019b) and Seurat (Hao et al, 2021). We treated genes that are differentially expressed as if they were co-clustered with the cells from the respective cluster (see Methods 4.13).

To be fair to all the methods, we tested numerous parameters choices for each method. For the summary statistics we ran every algorithm with 108 different parameter combinations that were chosen individually for each algorithm and are intended to represent the capabilities of the algorithm well. Crashed program runs and runs that resulted in a single bicluster encompassing the whole data were discarded.

Figure 6b shows that the overall clustering difficulty is higher for simulated data with a low number of differentially expressed genes and differential expression of smaller magnitude. However, the majority of biclustering algorithms struggle to cluster sparse scRNA-seq data irrespective of the simulated data (Fig. 6b). Besides CAbiNet, only Plaid and s4vd are able to generate meaningful biclusters. Although not being biclustering algorithms in the strict sense, both Seurat and Monocle3 perform well on the simulated data. In fact, Seurat shows the best performance of all tested algorithms on simulated data sets, while Monocle3 generally performs worse than CAbiNet. CAbiNet, however, outperforms all biclustering algorithms by a wide margin as well as Monocle3. Although the overall clustering error is lower for simulated data based on the pbmc3k data, s4vd seems to perform slightly better on this data set.

**Fig. 6.**
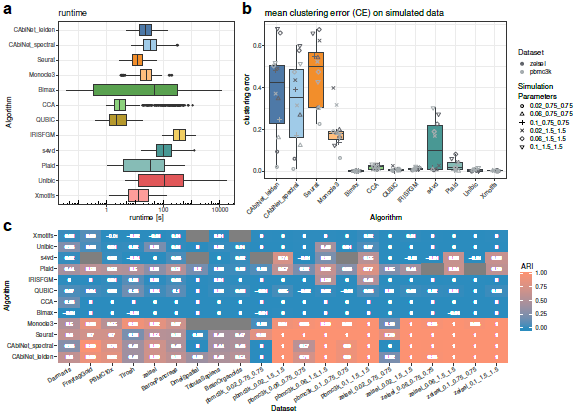
Benchmarking of CAbiNet biclustering against other biclustering algorithms and scRNA-seq analysis toolkits. **a**, Runtime across all parameter choices for simulated data. **b**, Mean clustering error (CE) of simulated data sets over all parameter choices. Different simulation parameters (the clustering “difficulty”) are differentiated by shape, whereas the data set used to estimate the base parameters (zeisel and pbmc3k) is marked by different shades of grey. **c**, Mean Adjusted Rand Index of (bi-)clustering results over all parameter choices for each algorithm on both real and simulated scRNA-seq data sets. The entries colored as grey indicate a failure of the respective algorithm on a certain data set. This can be due to either all runs failing on a data set or the algorithm only detecting a single bicluster.

Interestingly, CAbiNet is the only biclustering algorithm in the comparison that produces any meaningful results for the hardest simulations. Figure 6a further shows that CAbiNet runs similarly fast as other biclustering algorithms.

### 2.7 Evaluation on expert annotated data

To assess the performance of CAbiNet on experimental data we tested it on nine expert annotated scRNA-seq data sets. Again, CAbiNet gets compared to the same eight biclustering algorithms as above and to the clustering routines from Seurat and Monocle3. Since the expert clustering that comes with the data sets only provides us with a clustering of cells without genes, the ARI but not the CE is used as a quality measure.

The data sets range in size from 461 up to 35,291 cells and have been generated with different sequencing technologies (see Methods 4.12). Similarly to the benchmarking on simulated data, we again used 108 parameter combinations for each algorithm to ensure an even playing field.

Compared to the biclustering methods, CAbiNet with Leiden clustering on average yields the highest ARI of the cell clustering for all data sets except for one (Tirosh), where Plaid achieved the highest average ARI (Fig. 6c). CAbiNet with spectral clustering performs slightly worse than CAbiNet with Leiden for almost all data sets. However, it still outperforms other biclustering algorithms with the exception of Plaid on the ‘Darmanis’ and ‘Tirosh’ data sets. Some biclustering algorithms such as Xmotifs and s4vd completely failed in detecting any cell clusters for some data sets (marked as grey blocks in Fig. 6c), either because of issues in the algorithms’ implementation or because the entire data set was assigned to a single cluster.

Comparing with scRNA-seq cell clustering by Monocle3 and Seurat, CAb-iNet on average outperforms Monocle3 on all data sets. In 8/9 data sets CAbiNet produces a better cell clustering than Seurat, which only outperforms CAbiNet on the Darmanis data. Notably, Monocle3 fails to recognize cell clusters in the largest three data sets (Dmel, tabula sapiens and brain organoids) while CAbiNet still obtains reasonable clustering accuracy.

## 3 Discussion

We introduced CAbiNet and the biMAP as a novel visualization method to simultaneously plot cells and genes, allowing for interactive data exploration. The biMAP is a planar embedding placing genes with the cell clusters whose cells express the gene. It thus allows researchers to better understand cell-gene relationships and to annotate cell types easily.

We showed the applicability of CAbiNet to a wide variety of data sets and highlighted multiple ways in which CAbiNet can be used to generate novel insights. We demonstrated the general usage of CAbiNet to identify cell types on the PBMC10x and tabula muris data sets. Even in highly complex data sets, such as the brain organoids or spatial *Drosophila melanogaster* data that include developmental trajectories, CAbiNet is able to facilitate the differentiation of sub-types based on the 2D layout of cells and co-clustered marker genes. Moreover, CAbiNet is capable of refining the cell types in the spatial data and recognizes sub-types of epidermis and differentiates gastric caecum from midgut.

CAbiNet implements a novel biclustering approach. Benchmarking on the simulated data showed that CAbiNet is the best performing biclustering algorithm in comparison to a suite of other established algorithms. CAbiNet is overall the best tested biclustering algorithm on the experimental scRNA-seq data sets, where it outperforms all other algorithms on all but one data set. Comparing with the cell clustering algorithms Seurat and Monocle3, CAbiNet performs slightly better than Seurat on real data, but slightly worse than Seurat on simulated data sets. CAbiNet and Monocle3 have comparable performance on both simulated and small-sized real data sets, while Monocle3 failed on large-sized real data. In a real application without knowledge of the true clustering, the visual representation of genes and cell clusters in the biMAP provides additional information as to the reliability of a clustering and its marker genes.

An inherent limitation of CAbiNet is that it can only detect upregulated genes. The method is insensitive to genes which are characteristically down-regulated in a certain cell cluster. However, as cell types are generally defined through the expression and not the absence of specific genes, this is hardly an obstacle to cell type annotation. Another direction for future development would be to allow for a soft assignment of genes to clusters. This would provide a more realistic representation of genes that are involved in several cell types. Due to the fact that CAbiNet includes both cells and genes in the graph, the memory requirements are higher than in traditional cell-clustering approaches. Although we have ameliorated the problem by feature selection, there is probably still room to improve the implementation of CAbiNet.

CAbiNet is a valuable tool for researchers to gain a more intuitive understanding of their data and can be easily integrated into common workflows. The biMAP helps to visualize previously hidden cell-gene relationships in the data and thus enables the generation of new hypothesis. Complementary to commonly used methods, the biMAP can facilitate the verification of differential gene expression results and thereby support the definition of cell-types.

## 4 Methods

### 4.1 Data Pre-processing

Before applying CAbiNet to single-cell RNA-seq data, we recommend to pre-process the data. Removing unexpressed genes or cells with too many dropouts not only speeds up the computation but can also improve the clustering results. If not mentioned otherwise, we processed real and simulated scRNA-seq data as follows: First, outlier cells were filtered with the functions `perCellQCMetrics` and `perCellQCFilters` from the Bioconductor tool `scuttle` (McCarthy et al, 2017). They were then normalized by running `quickCluster`, `computeSumFactors` and finally `logNormCounts` from the Bioconductor package `scran` (Lun et al, 2016) with default parameters. We determined highly variable genes by first fitting a trend to the variance of the logcounts over all genes with respect to the mean expression with the function `modelGeneVar` from the package `scran`. The 80 % most variable genes with a biological component of the variance above 0 as determined by the function `getTopHVGs` were kept for further analysis. Additionally, all genes that are expressed in less than 1 % of all cells were discarded.

### 4.2 Correspondence Analysis

The pre-processed count matrix *X* with *n* cells and *m* genes is firstly transformed into a contingency table **P** by dividing each entry by *n*_++_, the overall sum of matrix entries.

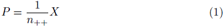

Each entry *p*_*ij*_ of **P** represents the observed probability a gene *i* is expressed in cell *j*. The row-sums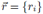 and column-sums 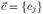 of **P** each add up to 1, where

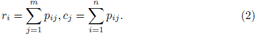

Thus one can define the expectation of an entry *e*_*ij*_ as the product of the i-th row-sum *r*_*i*_ and j-th column-sum *c*_*j*_ in **P**, that is *e*_*ij*_ = *r*_*i*_*c*_*j*_. Finally, one can calculate the matrix **S** of Pearson Residuals as

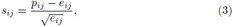

which describes the difference between observed and expected probabilities. Applying singular value decomposition the matrix *S* gets decomposed as

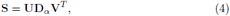

where **D**_*α*_ is a diagonal matrix with singular values *α*_*k*_ as the elements. The eigenvalue 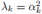 is defined as inertia in CA. Let 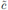 be the vector containing entries 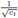 and 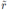 the vector made up of 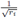. The standard coordinates of cells (**Γ**) and genes (**Φ**) can be obtained from the matrices **U** and **V**:

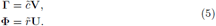

Scaling the standard coordinates by multiplying the respective singular values gives the principal coordinates of cells (**G**) and genes (**F**):

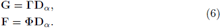

As a consequence, euclidean distances in the new space equal *χ*^2^-squared distances among the untransformed data. This property inspires us to use the principal coordinates to calculate gene similarities and cell similarities.

It is the hallmark of correspondence analysis that one can overlay these two spaces and merge the gene and cell data points. The technical prerequisite is that the gene points should be plotted in principal coordinates while the cells are plotted in standard coordinates. This ensures the geometric interpretation that when a gene-point and a cell-point lie far from the origin in the same direction, then they are highly associated. Intuitively, the association between samples and genes can be quantified by the association ratio, which is defined as the observed frequency divided by the expected frequency of an entry in a contingency table minus 1:

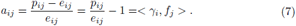

This formula implies that the association ratio between a gene and a cell equals the inner product of the respective gene-point (in principal coordinates) with the cell-point (in standard coordinates). This relationship geometrically connects cells with their marker genes. It allows us to build up the cell-gene graph and to cluster genes and cells simultaneously.

### 4.3 Dimension Reduction

In order to reduce the dimensionality of the data and to remove noisy, uninformative dimensions we reduce the data into *K* dimensions which preserve the most principal inertia. This is done through the function cacomp from the R Bioconductor package APL (Gralinska et al, 2022).

Due to the reduced dimensions, the association ratio between genes and cells would also change slightly. Using the standard and principal coordinates, the inner product of the row-points and column-points in the *K* dimensional reduced space can approximate the association ratio such that

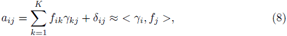

where *δ*_*ij*_ is an error term. Similarly, the Euclidean distance between principal coordinates in the dimensional reduced space approximates the *χ*^2^ distance between items in the original data.

#### 4.3.1 Gene ranking

In order to rank genes for each cluster we borrow the concept of S_*α*_-scores from Association Plots as described by Gralinska *et al*. (Gralinska et al, 2022). We computed the association plot coordinates (*x, y*) for genes in each cluster and then determined the angle *α* by randomly permuting the data to determine above which angle 99 % of genes lie due to chance in the Association Plot. The S_*α*_-score is then computed as:

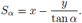

Co-clustered genes were then ranked by their S_*α*_-score.

### 4.4 Cell-Gene Graph

The cell-gene graph is constructed from four sub-graphs: a cell-cell graph, a gene-gene graph, a cell-gene graph and finally a gene-cell graph. Firstly, cell nodes are connected to the *k*_*c*_-nearest cell nodes by the euclidean distance, and to the *k*_*cg*_ gene nodes based with the highest Association Ratio in dimension reduced space.

In practice, genes that are not connected to any cell are removed as they are unlikely to be marker genes. If a high *k*_*cg*_ is chosen, there could also be genes erroneously connected with cells. CAbiNet provides users with an optional gene-pruning step to remove these genes: for each edge from a gene to a cell CAbiNet calculates how many of the cell’s direct neighbors also have an edge to this gene. If more than a user defined percentage of neighboring cells have an edge to the gene it is kept and removed otherwise.

The remaining genes are then connected to the *k*_*gg*_-closest gene nodes based on euclidean gene-gene distances. Finally, the gene-cell graph is obtained by simply transposing the adjacency matrix of the cell-gene graph.

The four sub-graphs are combined by joining their adjacency matrices to form a cell-gene graph. We further transform this nearest neighbor graph into an undirected, weighted shared-nearest-neighbor (SNN) graph. Shared Nearest Neighbors have been demonstrated to be a robust measure of similarity in high dimensions and to improve over the underlying primary measure such as euclidean distance (Houle et al, 2010). Edges in the SNN-graph are weighted by the Jaccard index (Tanimoto, 1958) between the neighborhoods of two nodes.

With the Jaccard index bounded between 0 (no overlap) and 1 (complete overlap), we remove an edge between two nodes if it is below a set cutoff. For our method we decided on a default pruning cutoff of 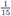.

### 4.5 biMAP Visualization

The SNN-graph also directly allows for the simultaneous visualization of cells and genes. CAbiNet converts the Jaccard similarities of the SNN-graph to a distance matrix and uses it as input to UMAP, which then produces what we call the biMAP. The biMAP embeds cells and genes in the plane as points of different size or shape to allow for quick distinction. For particularly crowded plots, cells can alternatively be summarized as hexagons, whose color will be mixed proportionally to the cluster representation. Similarly to a conventional UMAP, a biMAP can also be used to display the expression of a gene in the cells, but has the advantage to also show the location of the gene of interest in relation to all cells.

### 4.6 Biclustering

Many clustering algorithms can be applied to the SNN-graph and will automatically yield clusters containing both cells and genes. CAbiNet applies Leiden algorithm (Traag et al, 2019) as default clustering algorithm, and spectral clustering (Von Luxburg, 2007) as an alternative option.

### 4.7 PBMC10x data

The PBMC10x data set was pre-processed according to Methods 4.1 and the top 2,000 most variable genes were retained. This data matrix was subjected to SVD and 80 dimensions were kept using the Bioconductor package `APL`. The cell-gene graph was built up with `CAbiNet` with *k*_*c*_ = 20, *k*_*g*_ = 20, *k*_*cg*_ = 10 and *k*_*gc*_ = 50. Then Leiden clustering was applied to the graph to find biclusters. Getting the biclustering results from the function `caclust` in our package, we removed those clusters which contain fewer than 10 genes. The biMAP coordinates were calculated with the function `biMAP` with *k* = 10 and plotted with the function `plot_biMAP`. The feature biMAPs in Fig. 2c were drawn using `plot_feature_biMAP`.

### 4.8 Tabula Muris Limb Muscle data

The SmartSeq2 Tabula Muris data was downloaded through the R Bioconductor package `TabulaMurisData` (Soneson, 2021) and subsetted to the cells from the “Limb Muscle”. Unannotated cells were removed and the remaining cells were processed as outlined in Methods 4.1, with the exception of filtering for highly variable genes. We made sure that known marker genes do not get filtered out during the graph pruning, because we needed those genes for discussing the results.

We kept 40 dimensions after running correspondence analysis on the normalized log-counts and built the SNN-graph with *k*_*c*_, *k*_*cg*_ and *k*_*gc*_ set to 100 while we kept *k*_*g*_ to 50. The slightly lower *k*_*g*_ value for the gene-gene graph helps to reduce the agglomeration of genes away from cells and instead “pulls” them closer to the cells expressing the genes. In order to remove spurious edges, we pruned the kNN-graph with the parameter `overlap` set to 0.2. The biclustering was performed with Leiden and filtered for biclusters consisting of cells and genes.

### 4.9 Human cerebral organoids data

Data obtained from Rosebrock *et. al* (Rosebrock et al, 2022) was subsetted to organoids generated with the Triple-i protocol while keeping cells from all four cell lines. Unannotated cells as well as cells marked as doublets were excluded from the data. We kept all cells with a mitochondrial gene count of up to 40 %, as was done also in the original publication. Counts were normalized and pre-processed as described in Methods 4.1 and known marker genes are kept throughout the graph pruning.

Correspondence analysis was performed with the package `APL` and the first 80 dimensions were retained. The package `CAbiNet` was then used for the biclustering and joint visualization of cells and genes. The kNN sub-graphs were computed with `caclust` and the following parameters: *k* = 100 for the cell-cell and gene-gene sub-graphs and *k* = 75 for the cell-gene/gene-cell sub-graphs. The gene-cell graph was calculated as the transpose of the cell-gene graph and the genes were filtered by setting *overlap* = 0.2. Clustering was performed with the Leiden algorithm and biclusters that solely consisted of genes were removed. Cell cycle scores were computed with the scran function cyclone.

### 4.10 Spatial *Drosophila melanogaster* embryo data

The E14-16h *Drosophila melanogaster* embryo scRNA-seq data by Wang et. al (Wang et al, 2022) was pre-processed as described in Methods 4.1 and batch effects between spatial slices were removed with the ComBat (Johnson et al, 2006) function from the sva package (Leek JT, Johnson WE, Parker HS, Fertig EJ, Jaffe AE, Zhang Y, Storey JD, Torres LC, 2022). The data was then reduced to 150 dimensions by CA, and the cell-gene kNN graph was built up by using *k*_*c*_ = 60 for the cell-cell subgraph and *k* = 10 for gene-gene/cell-gene subgraphs. The gene-cell graph was set to the transpose of the cell-gene graph and genes were trimmed by graph pruning with *overlap* = 0.1. The resolution of `Leiden` was 1.2. For a clearer visualization of results, we trimmed out clusters with only genes or cells when plotting Fig. 5b-j.

### 4.11 Splatter simulated data

Single-cell RNA-seq data was simulated with the Bioconductor package `splatter` (Zappia et al, 2017), which allows the estimation of simulation parameters from real data. Parameters such as mean gene expression levels, library size, number of outliers or dropouts and the Biological Coefficient of Variation (BCV) are estimated from the given real data set and applied to the simulated data. For benchmarking we used two scRNA-seq data sets to estimate parameters: Zeisel Brain Data (Zeisel et al, 2015) (zeisel) and the PBMC3k data from 10x Genomics (10x Genomics, 2016) (pbmc3k). The zeisel data was downloaded through the R package `scRNAseq` Risso and Cole (2022) and the pbmc3k data through the R package `TENxPBMCData` Hansen et al (2022). For each simulation we simulated 1,000 cells, 10,000 genes and six clusters containing 25 %, 10 %, 10 %, 20 %, 30 % and 5 % of the cells respectively. For each set of estimated parameters six versions were made for which we varied the probability for a gene in a cluster to be differentially expressed as well as the shape of the log-normal distribution governing the magnitude of differential expression. The six combinations that were generated for each set of parameters as derived from the zeisel and pbmc3k data for a total of 12 datasets can be seen in Table 1.

**Table 1.**
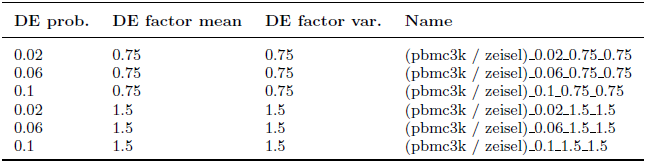
Simulated data sets. The six parameter combinations used to simulate scRNA-seq data based on either the zeisel or pbmc3k data sets. **DE prob**.: probability to be differentially expressed, **DE factor mean**: mean of log-normal distribution, **DE factor var**.: variance of log-normal distribution. **Simulated data sets**: name of 12 simulated data sets used in benchmarking.

| DE prob. | DE factor mean | DE factor var. | Name |
| --- | --- | --- | --- |
| 0.02 | 0.75 | 0.75 | (pbmc3k / zeisel)_0.02_0.75_0.75 |
| 0.06 | 0.75 | 0.75 | (pbmc3k / zeisel)_0.06_0.75_0.75 |
| 0.1 | 0.75 | 0.75 | (pbmc3k / zeisel)_0.1_0.75_0.75 |
| 0.02 | 1.5 | 1.5 | (pbmc3k / zeisel)_0.02_1.5_1.5 |
| 0.06 | 1.5 | 1.5 | (pbmc3k / zeisel)_0.06_1.5_1.5 |
| 0.1 | 1.5 | 1.5 | (pbmc3k / zeisel)_0.1_1.5_1.5 |

### 4.12 Experimental scRNA-seq data with ground truth cell types

Table 2 lists the scRNA-seq data sets with expert annotated or FAC sorted cell types. For a thorough comparison between CAbiNet and other algorithms, we included diverse data sets with different characteristics: The number of sequenced cells ranges from from 466 to 35,192 cells and the data sets have been produced through different scRNA sequencing methods such as 10x Genomics, Smart-seq2, Fluidigm C1, CEL-seq and Stereo-seq. Furthermore, they encompass data from different tissues such as brain, pancreas or blood as well as different organisms (mouse, human and *Drosophila melanogaster*). All these data sets were pre-processed as described in Methods 4.1.

**Table 2.** Experimental scRNA-seq data sets discussed in the results and used for benchmarking.

| Short Name | Dataset description | # Cells | # Genes | Protocol | Ref. |
| --- | --- | --- | --- | --- | --- |
| Darmanis | Human adult cortical samples | 466 | 22,085 | SMART-Seq2 | Darmanis et al (2015) |
| FreytagGold | Three human lung adenocarcinoma cell lines, HCC827, H1975 and H2228 | 925 | 58,302 | 10x | Freytag et al (2018) |
| tabula muris | Tabula Muris Limb Muscle | 1,960 | 23,433 | SMART-Seq2 | Schaum et al (2018) |
| zeisel | Mouse somatosensory cortex and hippocampal CA1 region (ZeiselBrain) | 2,874 | 14,508 | Fluidigm C1 | Zeisel et al (2015) |
| pbmc3k | Human peripheral blood mononuclear cells | 2,700 | 32,738 | 10x | 10x Genomics (2016) |
| Tirosh | Human melanoma tumor nonmalignant cells | 2,887 | 23,686 | SMART-Seq2 | Tirosh et al (2016) |
| PBMC10x | Human peripheral blood mononuclear cells (FACS sorted) | 3,362 | 33,694 | 10x | Ding et al (2020b) |
| BaronPancreas | Human Pancreas | 8,569 | 20,125 | CEL-seq | Baron et al (2016) |
| DemlSpatial | Drosophila melanogaster late stage embryo (14–16h after egg laying) | 15,295 | 13,668 | Stereo-seq | Wang et al (2022) |
| TabulaSapiens | Human endothelial cells | 32,701 | 58,559 | 10x | THE TABULA SAPIENS CONSORTIUM (2022) |
| BrainOrganoids | Human cerebral organoids | 35,291 | 33,538 | 10x | Rosebrock et al (2022) |

### 4.13 Benchmarking

Simulated data sets were created as explained in Methods 4.11. Both simulated and experimental data were pre-processed and normalized as described in Methods 4.1.

To allow for a fair comparison between (bi-)clustering algorithms we tested 108 parameter combinations for each algorithm and tried to spread them evenly over the reasonable parameter space. This number includes three choices for the number of highly variable genes used as input to the algorithms. We picked the top 2,000, 4,000 or 6,000 most highly variable genes similarly as during the pre-processing described in Methods 4.1. This can sometimes return a smaller number of genes than queried.

As some algorithms have more parameters than others, in practice this can mean, e.g. 18 variations of two parameters for one algorithm and only two variations of many parameters for another. Where possible we included the default parameter choices in the benchmarking. The exact parameters used for all the algorithms can be found in the scripts on GitHub (https://github.com/VingronLab/CAbiNet_paper). Runs that failed due to a tool’s implementation or inherent limitations were not rerun with other parameter choices.

### 4.14 Evaluation criteria

The similarity between detected and annotated ground truth cell clusters was calculated by the Adjusted Rand Index (ARI) (Hubert and Arabie, 1985), which ranges from 0 to 1. The larger the ARI the more detected clusters match with the ground truth. The similarity between detected and ground truth biclusters in simulated data is evaluated by the Clustering Error (CE) as defined by Horta and Campbello (Horta and Campello, 2014). The CE measures the proportion of matrix entries that are clustered differently after optimally matching the biclusters between the reference 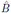 and the biclustering results *B* (Patrikainen and Meila, 2006):

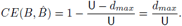

Here 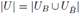 hwhere *U*_*B*_ is the union set of biclusters. *d*_*max*_ represents the maximal sum of overlapping bicluster elements between *B* and 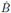. For our benchmarking we slightly modified the CE implementation from the python package biclustlib (Padilha and Campello, 2017).

## Acknowledgments

The authors wish to thank the IT group of the Max Planck Institute for Molecular Genetics for maintaining the computational resources in house and for their support. CK gratefully acknowledges financial support by the IMPRS-BAC PhD program. YZ gratefully acknowledges partial support by Shenzhen Science and Technology Program (Grant No. KQTD 20180411143432337, China). Y.H. and Q.H. acknowledge support by Shenzhen Key Laboratory of Gene Regulation and Systems Biology (Grant No. ZDSYS 20200811144002008, China) (Y.H.) and National Natural Science Foundation of China (Grant No. 32100684, Q.H.)

## Data availability

The simulated data sets we generated and the experimental data sets we used can be downloaded from Zenodo (https://zenodo.org/record/7433294#.Y6GJq6LMI10). Raw experimental data sets were sourced from the references listed in Table 2 and the supplementary material 7.

## Code availability

The codes to reproduce the results in this manuscript can be found on GitHub (https://github.com/VingronLab/CAbiNet_paper). The CAbiNet package can be installed from GitHub (https://github.com/VingronLab/CAbiNet).

## Supplementary information

Interactive biMAPs can be found on https://github.com/VingronLab/CAbiNet_paper/tree/main/SupplementaryMaterials.

---

**SFig. 1.**
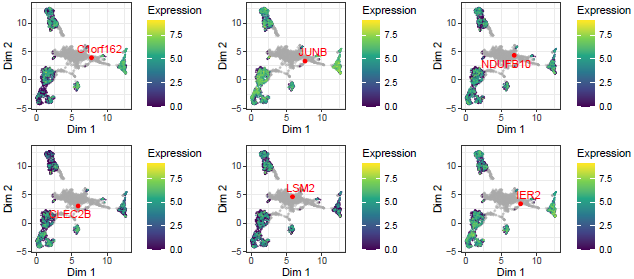
Feature biMAPs for PBMC10x data. The expression level of genes located in the center of the biMAP in Fig.2a. These genes are evenly expressed in more than one cluster, making them uninformative for defining cell types.

**SFig. 2.**
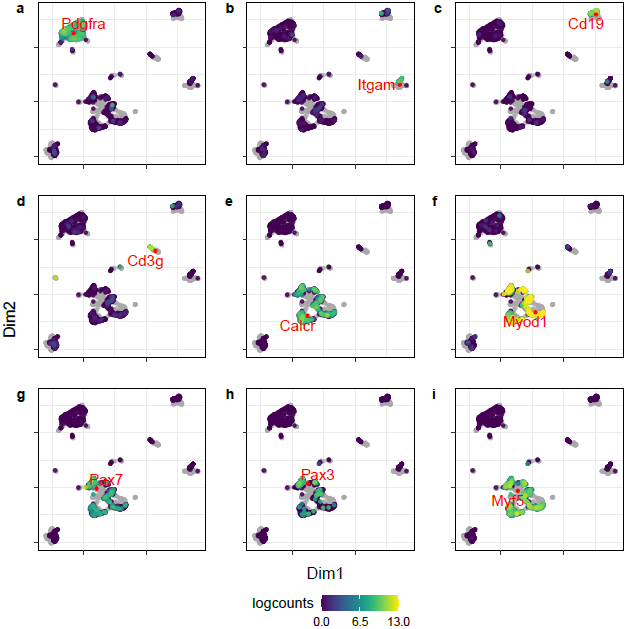
Feature biMAPs for Tabula Muris Limb Muscle data. **a-i**, biMAPs with cells colored by the log-expression of the genes mentioned in the text in Results 2.3.

**SFig. 3.**
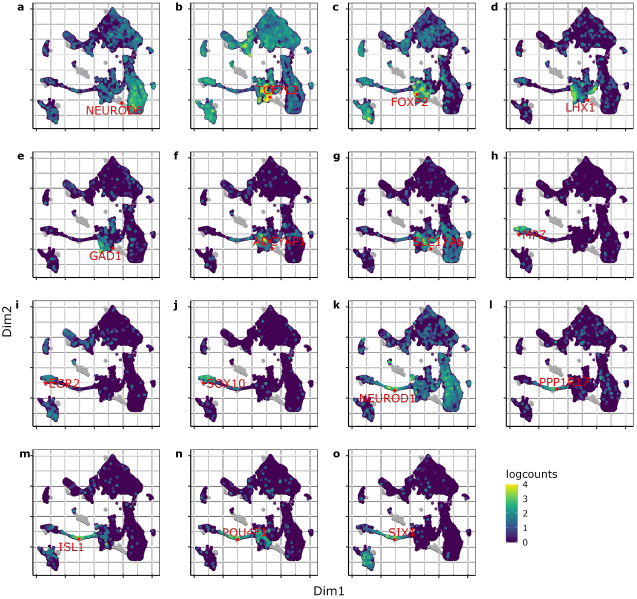
Feature biMAPs for the brain organoid data. **a-o**, biMAP with cells colored by the log-expression of the genes mentioned in the text in Results 2.4.

**SFig. 4.**
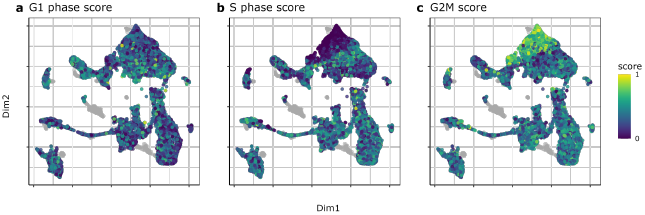
Cell cycle scores for the brain organoid data set. **a-c**, The biMAPs show the normalized cell cycles scores. The higher the score the likelier it is that the specific cell is in the respective phase of the cell cycle.

